# Training recurrent spiking neural networks in strong coupling regime

**DOI:** 10.1101/2020.06.26.173575

**Authors:** Christopher M. Kim, Carson C. Chow

## Abstract

Recurrent neural networks trained to perform complex tasks can provide insights into the dynamic mechanism that underlies computations performed by cortical circuits. However, due to a large number of unconstrained synaptic connections, the recurrent connectivity that emerges from network training may not be biologically plausible. Therefore, it remains unknown if and how biological neural circuits implement dynamic mechanisms proposed by the models. To narrow this gap, we developed a training scheme that, in addition to achieving learning goals, respects the structural and dynamic properties of a standard cortical circuit model, i.e., strongly coupled excitatory-inhibitory spiking neural networks. By preserving the strong mean excitatory and inhibitory coupling of initial networks, we found that most of trained synapses obeyed Dale’s law without additional constraints, exhibited large trial-to-trial spiking variability, and operated in inhibition-stabilized regime. We derived analytical estimates on how training and network parameters constrained the changes in mean synaptic strength during training. Our results demonstrate that training recurrent neural networks subject to strong coupling constraints can result in connectivity structure and dynamic regime relevant to cortical circuits.

## Introduction

With recent advances in machine learning, recurrent neural networks are being applied to a wide range of problems in neuroscience as a tool for discovering dynamic mechanisms underlying biological neural circuits. The activity of network models trained either on tasks that animals perform or to directly generate neural activity have been shown to be consistent with neural recordings. Examples include context-dependent computation (Mante et al., 2013), neural responses in motor cortex (Sussillo et al., 2015), decision-making with robust transient activities (Chaisangmongkon et al., 2017), and flexible timing by temporal scaling (Wang et al., 2018). However, due to the large number of parameters, the network connectivities are often not constrained to be biologically plausible, thus it remains unclear if and how biological neural circuits could implement the proposed dynamic mechanisms.

One approach for imposing biological constraints is to develop model architectures based on detailed connectivity structure of the neural system. The complete connectome is available for certain invertebrates such as Drosophila, and it is possible to build recurrent neural network models constrained by the detailed structure of the connectome (Litwin-Kumar and Turaga, 2019; Eschbach et al., 2020; Zarin et al., 2019). In cortical regions of higher animals, however, obtaining a connectome and applying it for computational modeling purposes pose significant technical and conceptual challenges. Therefore, it is important to develop alternative methods for constraining the connectivity in a biologically plausible manner independently of the detailed connectome structure.

Recent studies have applied training methods to rate-based and spiking cortical circuits of excitatory and inhibitory neurons (Song et al., 2016; Nicola and Clopath, 2017; DePasquale et al., 2016; Kim et al., 2019) but the focus was to preserve the excitatory-inhibitory structure (i.e. Dale’s law) without constraining the synaptic strength to a strong coupling regime (but see Ingrosso and Abbott (2019); Baker et al. (2019)) that exhibits chaotic dynamics (Sompolinsky et al., 1988), asynchronous spiking (van Vreeswijk and Sompolinsky, 1996; Renart et al., 2010; Rosenbaum et al., 2017), spiking rate fluctuations (Ostojic, 2014), and inhibition-stabilization (Tsodyks et al., 1997) as observed in neural recordings. Here, we show that by preserving the mean excitatory and inhibitory synaptic weights of a strongly coupled network, trained networks can exhibit large trial-to-trial variability in spiking activities, respect Dale’s law in most synapses without imposing strict constraints, and show a paradoxical phenomenon found in inhibition-stabilized networks. We provided analytical estimates on how the network and training parameters constrain the size of synaptic updates during network training. Our findings demonstrate that including synaptic constraints that maintain strong coupling strength allows spiking networks to respect dynamic properties of standard cortical circuit models in addition to achieving the learning goal.

## Results

### 1 Learning under excitatory and inhibitory synaptic constraints

The initial network consisted of an equal number of excitatory and inhibitory leaky integrate-and-fire neurons (*N*_*E*_, *N*_*I*_ = 500) randomly connected with connection probability *p* = 0.1. The recurrent synaptic weights *W*_*ij*_ and the external inputs were sufficiently strong such that the total excitatory inputs to neurons, i.e. the sum of recurrent excitatory current and external input, exceeded spike threshold, and inhibitory feedback was necessary to prevent run-away excitation (Fig. 1A.i). In this parameter regime, neurons emitted spikes asynchronously and irregularly (Fig. 1A.ii), and had large trial-to-trial variability as measured by the Fano factor of spike counts in a 500 ms time window across trials starting at random initial conditions (Fig. 1A.iii). The synaptic connections to every neuron were statistically identical in that the presynaptic neurons were randomly selected, but the number of excitatory (*pN*_*E*_) and inhibitory (*pN*_*I*_) synaptic connections were fixed and their weights were constant depending only the the connections type, *W*_*αβ*_, *α, β* ∈ {*E, I*} (i.e., fixed indegree and constant weights). This network configuration resulted in a homogeneous firing rate distribution (Fig. 1A.iv). Initial networks with non-identical connectivity statistics that generate wide rate distributions were also investigated in Section. 6.

**Figure 1:**
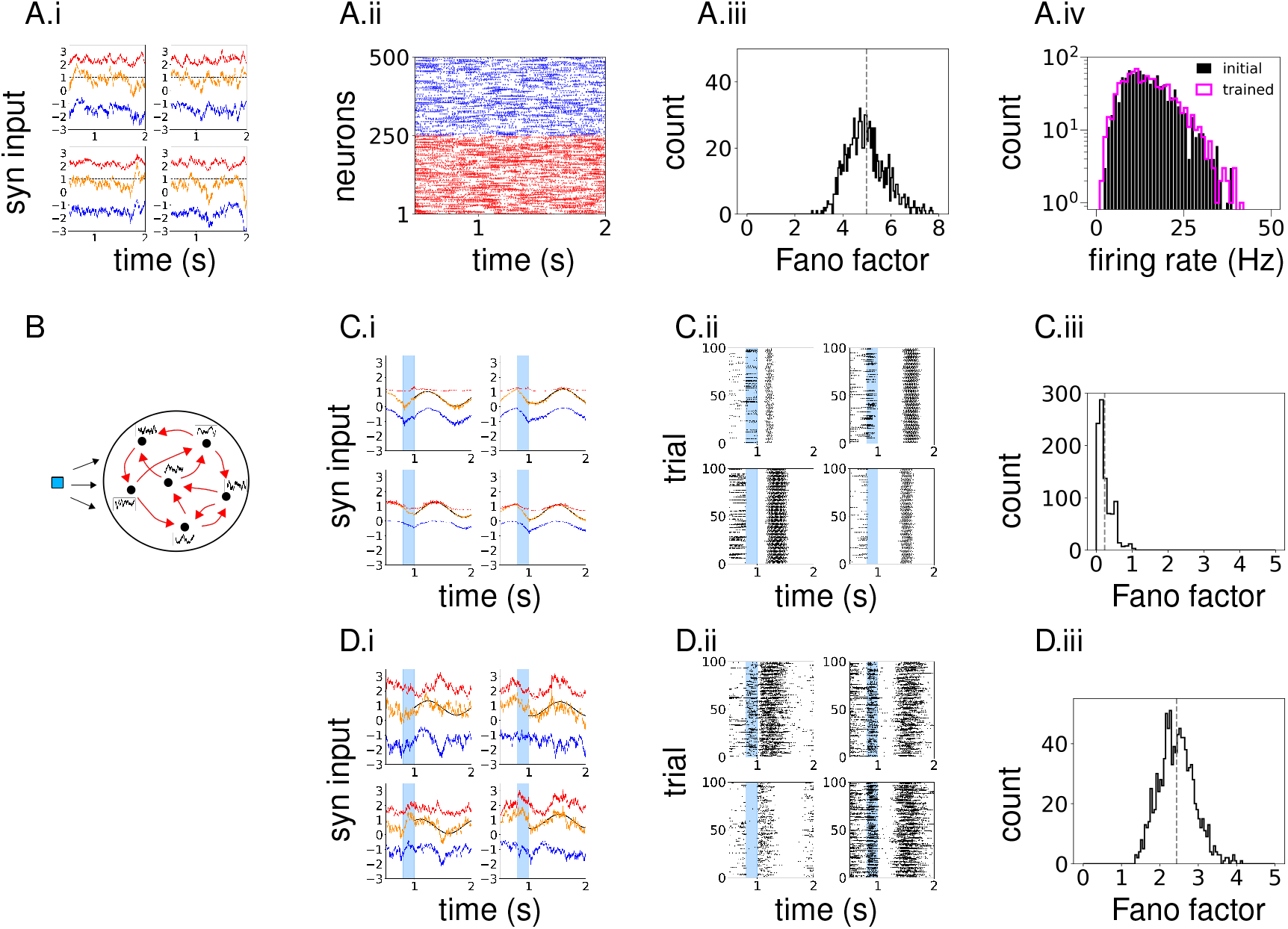
Comparison of FORCE- and ROWSUM-trained network activities. **(A)** Population activity of the initial network. (A.i) Synaptic inputs to example neurons; excitatory (red), inhibitory (blue) and the total (orange) inputs; external input is added to the excitatory and total inputs; dashed line indicates the spike threshold. (A.ii) Spike raster of sample neurons. (A.iii) Fano factor of all neurons; spike counts across trials in a 500 ms time window are used to compute the Fano factor and averaged over the training window to obtain a time-averaged Fano factor for each neuron. (A.iv) Firing rate distribution of neurons; initial network (black), ROWSUM-trained network (magenta). **(B)** Schematic of network training. Brief external input (blue) is applied to trigger learned response; individual neurons learn their own target patterns (black); recurrent synaptic connections (red) are modified to generate the target patterns. **(C)** Population activity of a FORCE-trained network. (C.i) Synaptic inputs to example neurons. (C.ii) Spike trains of the same neurons in (C.i) across trials starting at random initial conditions. (C.iii) Fano factor of trained neurons. **(D)** Population activity of a ROWSUM-trained network; Same color codes as in (C).

In the present study, spiking neural networks were trained to generate desired recurrent activity patterns. In related work, Rajan et al. (2016) trained a recurrent rate network to generate sequential activity observed in the posterior parietal cortex in mice, Laje and Buonomano (2013) stabilized inherent chaotic trajectories of the initial rate network, and Kim and Chow (2018) generated arbitrarily complex activity patterns in recurrent spiking networks. We point out that these tasks are different from the standard machine learning tasks, such as image classification, in that individual neurons within the network learn to generate certain activity patterns. Other studies have investigated performing machine learning tasks in spiking networks (Nicola and Clopath, 2017; Huh and Sejnowski, 2018).

Specifically, our goal was to train the synaptic current *u*_*i*_(*t*) to each neuron *i* = 1,…, *N* such that it followed target activity pattern *f*_*i*_(*t*) defined on time interval *t* ∈ [0*, T*]. To trigger the target response, each neuron was stimulated by a constant input with random amplitude for 200 ms (Fig. 1B). We treated every neuron’s synaptic current as a read-out, which made our task equivalent to training *N* recurrently connected read-outs. For spiking network models with current-based synapses, neuron *i*’s synaptic current *u*_*i*_ can be expressed in terms of the spiking activities of other neurons *r*_*j*_, *j* = 1,…, *N* through the exact relationship *u*_*i*_ = Σ_*j*_ *W*_*ij*_*r*_*j*_ (see Eqs. 11 and 14 for details). Therefore, we adjusted the incoming synaptic connections *W*_*ij*_, *j* = 1,…,*N* to neuron *i* by the recursive least squares algorithm (Haykin, 1996) in order to generate the target activity. This training scheme allowed us to set up independent objective functions for each neuron and potentially update them in parallel.

The objective function included the error between the synaptic currents and target patterns, and, importantly, we considered two types of synaptic regularization The first type regularized the standard *L*_2_-norm of the incoming synaptic weights to each neuron *i*,

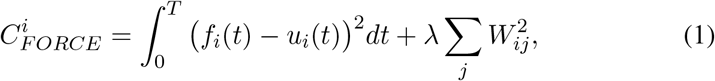

and 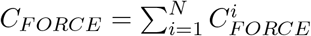 was the full objective function. The second type regularized the sums of excitatory and inhibitory synaptic weights, respectively, to each neuron *i*, in addition to the *L*_2_-norm,

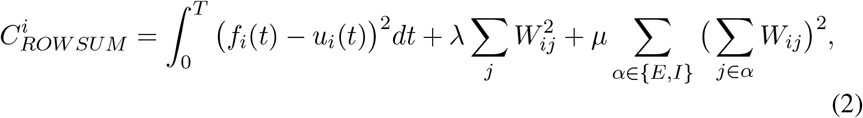

and 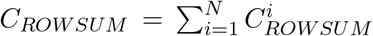 was the full objective function. The first type, which is equivalent to the algorithm of Sussillo and Abbott (Sussillo and Abbott, 2009) and used previously in spiking networks (Kim and Chow, 2018; Nicola and Clopath, 2017), will be referred to as FORCE training. The second type will be referred to as ROWSUM training since it regularizes the sum of excitatory and inhibitory weights, respectively, in each row of the connectivity matrix.

We note how the hyperparameters λ and *μ* can be interpreted. At first glance, it appears that large positive values of λ and *μ* should reduce the magnitude of synaptic weights in trained networks. However, the recursive least squares is different from the standard gradient descent algorithms in that the synaptic update rule is not derived from the gradient of cost function. Instead, it finds a critical point to the difference of gradients of cost function at two consecutive time points (see Methods / Section 2). This implies that large positive values of λ and *μ* penalize the size of synaptic updates, resulting in small synaptic updates or slow learning rate (see Sussillo and Abbott (2009) for further discussion). Therefore, the synaptic weights trained with large λ and *μ* stay close to their initial weights if trained by the recursive least squares algorithm.

We first considered simple target patterns consisting of sinusoidal waves 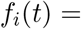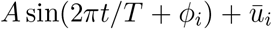 defined on t ∈ [0, *T*] with duration *T* = 1000 ms, random phase *ϕ*_*i*_, fixed amplitude *A* and bias 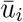. The amplitude *A* was the standard deviation of synaptic currents in time averaged over all neurons, and the bias 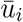 for neuron *i* was the mean synaptic current to neuron *i* in the initial network. After adjusting the recurrent weights with two training schemes, we examined the differences in trained network activities. Notably, the large fluctuations of excitatory and inhibitory synaptic currents in the initial network were not observed in FORCE-trained networks. Instead, the total synaptic currents closely followed the target patterns with little error (Fig. 1C.i). Such synaptic currents led to highly regular spikes with little variability across trials starting at random initial conditions (Fig. 1C.ii), resulting in a small Fano factor (Fig. 1C.iii). In contrast, ROWSUM-trained networks, which were encouraged to preserve the initial excitatory and inhibitory synaptic weights, exhibited strong synaptic fluctuations, irregular spike trains and large trial-to-trial variability (Fig. 1D.i,ii,iii). Also, the firing rate distribution of trained neurons closely followed the firing rate distribution of untrained neurons, showing that sinusoidal targets minimally altered the firing rate distribution (Fig. 1A.iv).

### 2 ROWSUM constraint can preserve the mean excitatory and inhibitory synaptic strength

We examined the trained synaptic weights to better understand the underlying network structure that led to different levels of variability in neuron activities in the FORCE- and ROWSUM-trained networks. We found that the excitatory and inhibitory synaptic weights of the initial network were both redistributed around zero by the FORCE learning (Fig. 2A.i), and the fraction of synapses that violated Dale’s law increased gradually as the number of training iteration increased (Fig. 2A.ii). The mean strength of excitatory and inhibitory synaptic connections to each neuron also moved towards zero (Fig. 2A.iii). This shift in synaptic weight distributions can be explained by the *L*_2_-regularization term in Eq. 1 that allowed small synaptic weights, and as a consequence, the mean excitatory and inhibitory synaptic connections to each neuron were significantly smaller than those of the initial network. After removing synaptic connections that violated Dale’s law, the FORCE-trained network was able to generate the target patterns with high accuracy (Fig. 2A.iv). These findings suggested that the network connectivity resulting from FORCE training was degenerating and highly tuned to produce the target dynamics with minimal error.

**Figure 2:**
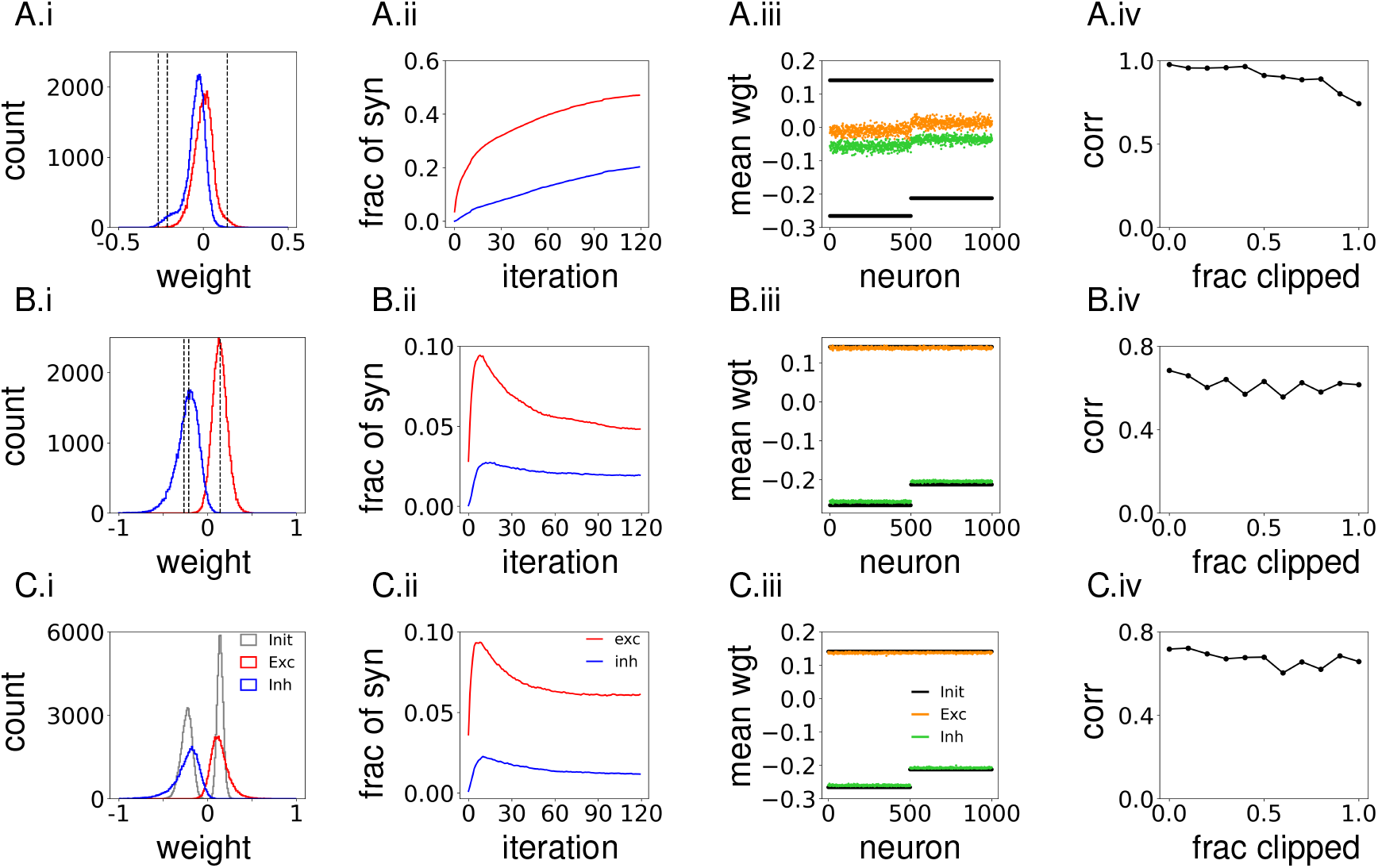
Network connectivity of trained networks. **(A)** FORCE-trained network. (A.i) Distribution of synaptic weights of the FORCE-trained network (120 training iterations); red (blue) shows synapses that are excitatory (inhibitory) in the initial network; dotted line shows the initial synaptic strength. (A.ii) Fraction of excitatory (red) and inhibitory (blue) synapses that violate Dale’s law after each training iteration. (A.iii) Mean strength of synaptic connections to individual neurons after FORCE-training; orange (green) dots show connections that are excitatory (inhibitory) in the initial network; black dots show the initial mean synaptic weights. Mean excitatory and inhibitory weights to neurons were identical in the initial network since indegrees were fixed and synaptic weights were constant. (A.iv) Performance of trained networks, measured as the correlation between target patterns and synaptic activities, when synapses violating Dale’s law are removed; fraction of clipped synapses equals 1 if all synapses violating Dale’s are removed. **(B)** Same as in (A), but for a ROWSUM-trained network with fixed indegree and constant weights. **(C)** Same as in (A), but for a ROWSUM-trained network with random indegrees and Gaussian weights; correction terms are added to reconcile differences in synaptic weights each neuron receive.

For the ROWSUM training, we used the same network and training parameters as the FORCE training but included the additional penalty that constrained the sums of excitatory and inhibitory synaptic weights, respectively, to each neuron. We found that the excitatory and inhibitory weights of the trained synapses were distributed around their initial values (Fig. 2B.i), and the fraction synapses that violated Dale’s law increased in the early stage of training but gradually decreased and stabilized as the training continued (Fig. 2B.ii). The mean strength of excitatory and inhibitory synaptic connections to each neuron after training deviated little from the initial mean synaptic weights (Fig. 2B.iii), demonstrating that the ROWSUM-regularization allowed the trained network to preserve the mean excitatory and inhibitory strength of the strongly coupled initial network. The fraction of synapses that violated Dale’s law after training (approximately 5% and 2% of excitatory and inhibitory connections, respectively) was significantly smaller than a FORCE-trained network (40% and 20%, respectively), and removing them had only a small impact on the network performance (Fig. 2B.iv).

Next, we investigated if ROWSUM training was applicable to initial networks with random indegrees and weights. To this end, we constructed an Erdos-Renyi graph where each synaptic connection was generated independently with probability *p*, i.e., Pr(*W*_*ij*_) = *p* if neuron *j* connects to neuron *i*, resulting in a connectivity matrix with a non-constant indegree distribution. The synaptic weight of each connection was sampled from Gaussian distribution with mean *W*_*αβ*_ and standard deviation *W*_*αβ*_/5 for *α, β* ∈ {*E, I*} to ensure the initial network respected Dale’s law. In order to reconcile differences in synaptic weights neurons received, we added a correction term to nonzero elements in each row of the connectivity matrix, such that the sum of excitatory and inhibitory synaptic weights to neuron *i* ∈ *α* was equal to *pN*_*E*_*w*_*αE*_ and *pN*_*I*_*w*_*αI*_, respectively. Including such correction terms resulted in a homogeneous rate distribution, as opposed to a wide rate distribution, and trained networks qualitatively agreed with those initialized with fixed indegrees and constant weights (Fig. 2C.i-iv).

These findings demonstrated that constraining the sums of excitatory and inhibitory synaptic weights, respectively, allowed the individual synapses to learn target patterns while keeping the mean synaptic strength close to the initial values. Since the excitatory and inhibitory synaptic weights of the initial network were strong enough to generate variable spiking activity, the trained network was able to inherit the dynamic property of the initial network.

Before turning to limitations of training complex target patterns and initial networks without the correction terms in Sections. 4 and 6, we examined if other aspects of the initial network dynamics were inherited by the trained network when the mean excitatory and inhibitory synaptic weights were preserved.

### 3 ROWSUM constrained networks inherit dynamic features of inhibition-stabilized network

The initial network was set up to be an inhibition-stabilized network (ISN), where strong recurrent excitatory connections were stabilized by inhibitory feedback without which the network would exhibit run-away excitation (Tsodyks et al., 1997; Sanzeni et al., 2019; Sadeh and Clopath, 2020). An ISN exhibits a paradoxical phenomenon where an external stimulus that excites inhibitory neurons decreases their firing rates. Although the stimulus can transiently increase the firing rates of inhibitory neurons, this activity inhibits excitatory neurons that in turn reduce recurrent excitatory input to inhibitory neurons. The reduction in recurrent excitation is larger than the external stimulus resulting in a net decrease in excitatory input to inhibitory neurons. Such large reduction in recurrent excitation occurs because strongly connected excitatory neurons respond to changes in input with large gain (Tsodyks et al., 1997; Sanzeni et al., 2019; Sadeh and Clopath, 2020).

To verify that the initial network operated in the inhibition-stabilized regime, we injected additional excitatory stimulus to inhibitory neurons (Fig. 3A) and found that both excitatory and inhibitory population firing rates decreased (shaded region in Fig. 3B.i). As the external stimulus to inhibitory neurons was further increased, the excitatory population rate decreased gradually, and once the excitatory population activity was fully suppressed the firing rate of inhibitory neurons started increasing with the external stimulus (Fig. 3B.ii). These dynamic features are consistent with theoretical analysis of ISNs that the paradoxical phenomenon occurs only within a limited range of stimulus intensity. In a strong stimulus regime, the network is effectively reduced to a single inhibitory population, therefore the paradoxical phenomenon can no longer exist.

**Figure 3:**
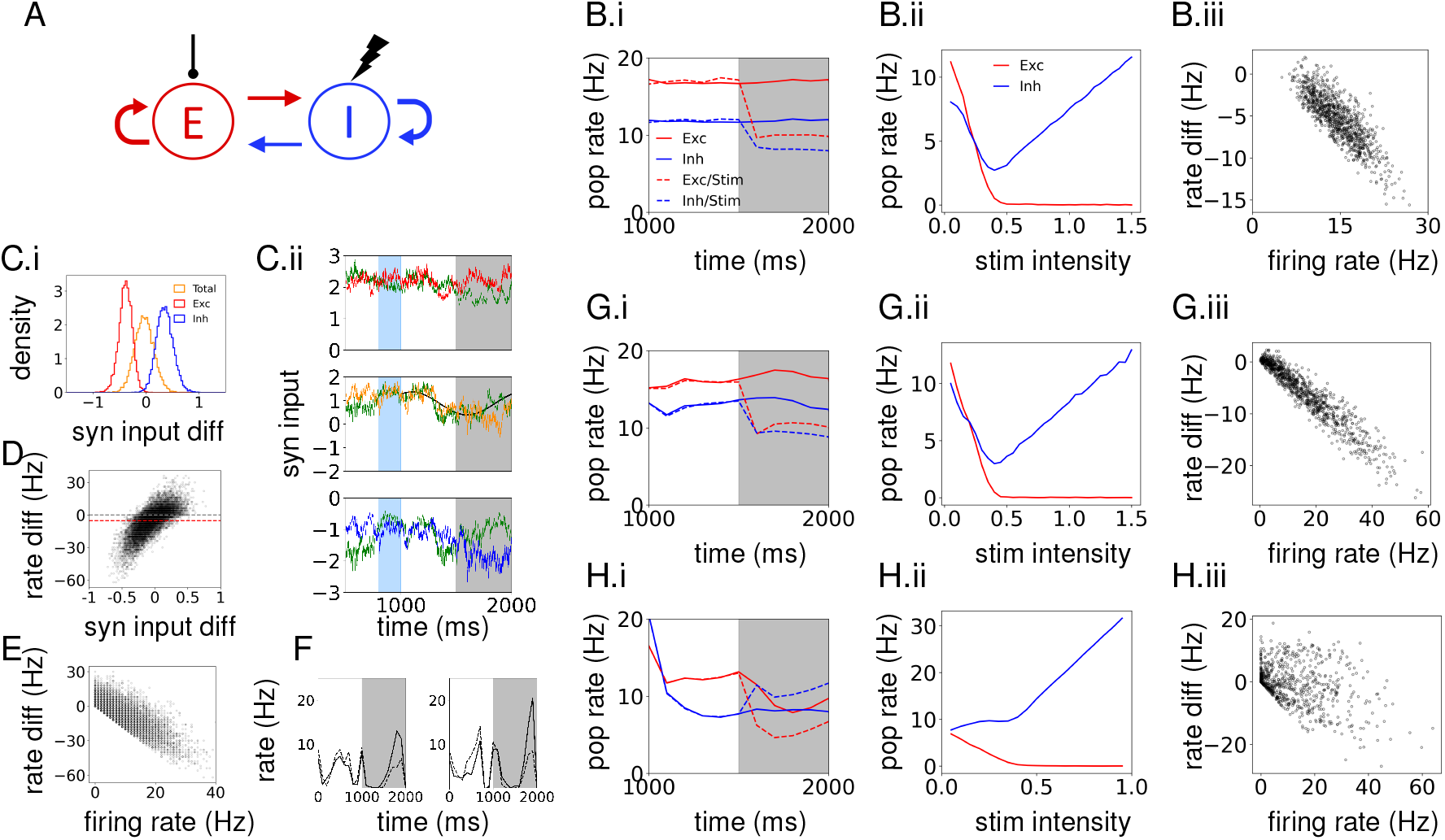
ROWSUM-trained networks exhibit dynamic features of inhibition-stabilization. Shaded region shows the time period during which inhibitory neurons are excited by additional external stimulus. **(A)** Schematic of network structure where inhibitory neurons are stimulated. **(B)** Initial network activities in response to stimulating inhibitory neurons. (B.i) Excitatory (red) and inhibitory (blue) population firing rates with (dotted) and without (solid) the additional stimulus to inhibitory neurons. (B.ii) Population firing rates as a function of stimulus intensity. (B.iii) Trial-averaged firing rates of individual neurons without stimulus vs. changes in neurons firing rates due to the stimulus. **(C)** Synaptic inputs to individual neurons. (C.i) Distribution of changes in time-averaged synaptic inputs to individual neurons due to the stimulus. For excitatory (inhibitory) inputs, negative (positive) value implies reduction in the average input; excitatory (red), inhibitory (blue) and total (orange). (C.ii) Synaptic inputs to a neuron in an unstimulated network is compared to those in a stimulated network (green). **(D)** Changes in total synaptic input vs. changes in the number of emitted spikes due to the stimulus; dots show individual neurons. **(E)** The number of emitted spikes in an unstimulated network vs. changes in the number of emitted spikes due to the stimulus; dots show individual neurons. **(F)** Trial-averaged firing rate of sample neurons with (dotted) and without (solid) the stimulus. **(G)** Same as in (B), but for a ROWSUM-trained network. **(H)** Same as in (B), but for a FORCE-trained network.

To better understand how individual neurons responded to the external stimulus, we stimulated inhibitory neurons over multiple trials and examined the trial-averaged firing rates of individual neurons with and without the stimulus. The relationship between the trial-averaged firing rate of unstimulated neurons and changes in their firing rates due to the stimulus revealed that neurons with high baseline firing rates resulted in large reduction in their spiking activities (Fig. 3B.iii).

Since the ROWSUM training preserved the mean synaptic strength of the initial network, we investigated if, in addition to learning target activity patterns, the trained network preserved unique features of initial network’s population dynamics characterized by the strong excitation and feedback inhibition. To test if the ROWSUM-trained network exhibited the dynamic features of ISN, we evoked the learned activity patterns in the trained network, and then injected external stimulus that excited all inhibitory neurons during the second half of the target patterns. We found that, although the temporal average of recurrent excitatory and inhibitory inputs to individual neurons were both reduced in their magnitudes due to the stimulus (Fig. 4Ci, red and blue), the distribution of total synaptic inputs to neurons were centered around zero and did not exhibit apparent hyperpolarizing effects at the population level (Fig. 4Ci, orange). When we examined single trial responses with and without the stimulus, neurons whose total synaptic inputs were hyperpolarized by the stimulus (negative side of the orange curve in Fig. 4C.i) had large reduction in their spiking activities (negative side in Fig. 4D) in comparison to the increased spiking activities (positive side in Fig. 4D) in other neurons whose total synaptic inputs were depolarized by the stimulus (positive side of orange curve in Fig. 4C.i). As a result, the population firing rate decreased by 5 Hz (red line in Fig. 4D) consistently with the firing rate reduction seen in the stimulated initial network (Fig. 4B.i). In addition, neurons with high spiking rate in the unstimulated network had large reduction in their emitted spikes (Fig. 3E) in agreement with the trial-averaged response in the initial network (Fig. 3B.iii).

**Figure 4:**
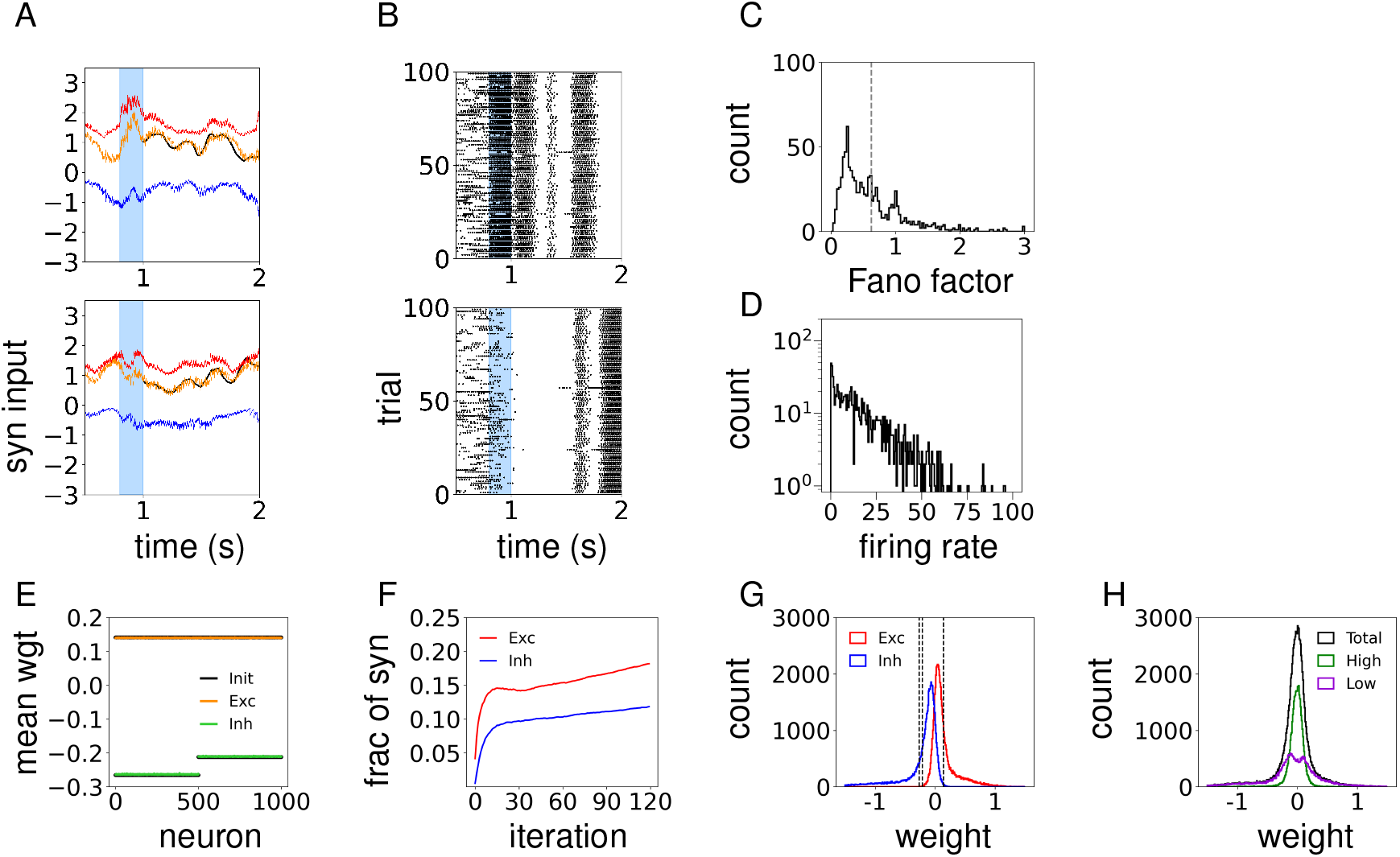
Learning complex activity patterns under the ROWSUM constraint on excitatory and inhibitory synapses. Target patterns consist of trajectories generated from Ornstein-Ulenbeck process. **(A)**Synaptic inputs to trained neurons; excitatory (red), inhibitory (blue) and the total (orange) inputs; external input is added to the excitatory and total inputs. **(B)**Spike trains of the same neurons in (A) across trials. **(C)**Fano factor of trained neurons; dashed line indicates the mean. **(D)**Distribution of neurons firing rates during the trained time window. **(E)**Mean strength of trained synaptic weights to each neuron; orange (green) dots show the mean strength of synaptic weights that are excitatory (inhibitory) prior to training; black dots show the mean strength of initial excitatory (positive) and inhibitory (negative) synaptic weights. **(F)**Fraction of excitatory (red) and inhibitory (blue) synapses that violate Dale’s law after each training iteration. **(G)**Distribution of trained synaptic weights; red (blue) shows synapses that are excitatory (inhibitory) in the initial network; dotted line shows the initial synaptic strength. **(H)**Distribution of trained synaptic weights; all the synapses (black); outgoing synapses from top (green) and bottom (purple) 40% of neurons ordered by their firing rates.

The effects of exciting inhibitory neurons were observed more clearly in trial-averaged firing rates in individual neurons and population rates. Without additional stimulus to inhibitory neurons, the trial-averaged firing rates followed the target rate patterns as trained (Fig. 3F). However, when the external stimulus was injected to excite the inhibitory neurons, their firing rates decreased compared to those of unstimulated neurons and similar effects was observed in both the excitatory and inhibitory population firing rates (Fig. 3G.i). We also found that the paradoxical effect was observed in a limited stimulus range for the ROWSUM-trained network (Fig. 3G.ii), and the neurons with high trial-averaged firing rates in an unstimulated network had larger reduction in their spiking activities due to the stimulus (Fig. 3G.iii). These findings show that the ROWSUM-trained network inherited the dynamic features of the initial network.

To test if the FORCE-trained network exhibited similar dynamic features, we trained an inhibition-stabilized network without the ROWSUM regularization term. Injecting excitatory inputs to inhibitory neurons in the FORCE-trained network did not elicit the paradoxical effect: the inhibitory population rate increased and the excitatory population rate decreased (Fig. 3H.i). Moreover, the paradoxical effect was not observed over a wide range of stimulus intensity (Fig. 3H.ii). Also, a large number of neurons increased their trial-averaged firing rates due to the stimulus (Fig. 3H.iii). These findings suggest that the FORCE-trained network did not operate in the inhibition-stabilized regime.

### 4 Limitations of constraining excitatory and inhibitory synaptic weights

So far, (1) the initial network connectivity was adjusted such that the sum of excitatory and inhibitory synaptic weights to neurons were uniform, and (2) the target patterns were simple sinusoidal perturbations of the initial synaptic activities. Such initial network setup and target patterns led to relatively homogeneous firing rate distribution. We asked if the ROWSUM training was still effective when either the initial network connectivity or the set of target patterns was no longer fine-tuned. To this end, we investigated if both the initial network setup and the choice of target functions can limit the effectiveness of ROWSUM training. In this section, we considered initial networks whose total excitatory and inhibitory synaptic weights across neurons were uniform as before, but neurons were trained on complex target patterns to encourage a wide range of activity patterns. Initial networks with non-identical synaptic weights across neurons is discussed in Section. 6.

To generate complex target patterns we sampled trajectories from Ornstein-Ulenbeck process with decay time constant 200 ms and applied the moving-average with 100 ms time window to smooth out fast fluctuations. The amplitude of trajectories was scaled so that their standard deviation matched with that of synaptic currents of initial network. Each neuron was trained on independently generated trajectories with the ROWSUM constraints on the sum of excitatory and inhibitory synaptic weights, respectively. We found that, although the trained neurons successfully tracked the complex trajectory patterns (Fig. 4A), the trial-to-trial variability was relatively low (Figs. 4B, C) and the firing rate distribution was wide including many low firing rate neurons (Fig. 4D).

Examining the trained weights showed that the mean strength of excitatory and inhibitory synaptic connections to each neuron did not deviate from their initial values, showing that the ROWSUM constraints worked effectively (Fig. 4E). However, the fraction of synapses that violated Dale’s law increased gradually with training iterations (Fig. 4F), and the distribution of synaptic weights of the trained network showed that a large number of synapses were degenerating (Fig. 4G). In particular, the synapses outgoing from neurons with high firing rate were concentrated near zero, in contrast to those from neurons with low firing rates (Fig. 4H).

To better understand how the trained synaptic weights might depend on the firing rate distribution of presynaptic neurons, we analytically estimated the size of synaptic weight updates assuming presynaptic neurons followed simplified rate distributions (see Methods / Section 4 for details). The analytical expressions suggested that, for a homogeneous rate distribution, i.e., all neurons have the same rate, the size of synaptic update is small, i.e., *O*(1/(*μK*)), due to the cancellation effect by the ROWSUM-regularization (Eq. 31). On the other hand, for an inhomogeneous rate distribution consisting of high and low firing rate neurons, the size of synaptic update is large, i.e., *O*(1), and the synaptic connections from two neuron types are updated in opposite directions (Eq. 32). The analytical results from the inhomogeneous rate distribution were consistent with the full ROWSUM-training in that the synapses from high firing rate neurons concentrated near zero while those from low firing rate neurons had a wide distribution including strong synapses (Fig. 4H). Moreover, analysis of the homogeneous rate distribution suggested that applying the ROWSUM constraints to subpopulations that share similar activity levels can reduce deviations of individual synaptic weights from their mean values.

### 5 ROWSUM constraints on multiple subpopulations

In order to overcome the limitations of applying ROWSUM constraints to a population of neurons with a wide firing rate distribution, we further divided the neurons into multiple subpopulations where neurons within a subpopulatioin shared similar level of spiking activities. First, to estimate individual neuron’s firing rate after training, external inputs that followed the target patterns were injected to each neuron. Then, the excitatory and inhibitory neurons were respectively ordered by their time-averaged firing rates and divided into *M* subpopulations of equal sizes by assigning the *α*^*th*^ *N*_*E*_/*M* excitatory and *N*_*I*_/*M* inhibitory neurons to *E*_*α*_ and *I*_*α*_, respectively. Finally, the ROWSUM constraint was applied to synapses from each subpopulation *E*_*α*_, *I*_*α*_, *α* = 1,…, *M*, separately, resulting in the following synaptic constraint for training neuron *i*:

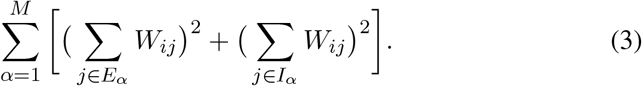

See Figure 5A for a schematic of constraining multiple subpopulations.

**Figure 5:**
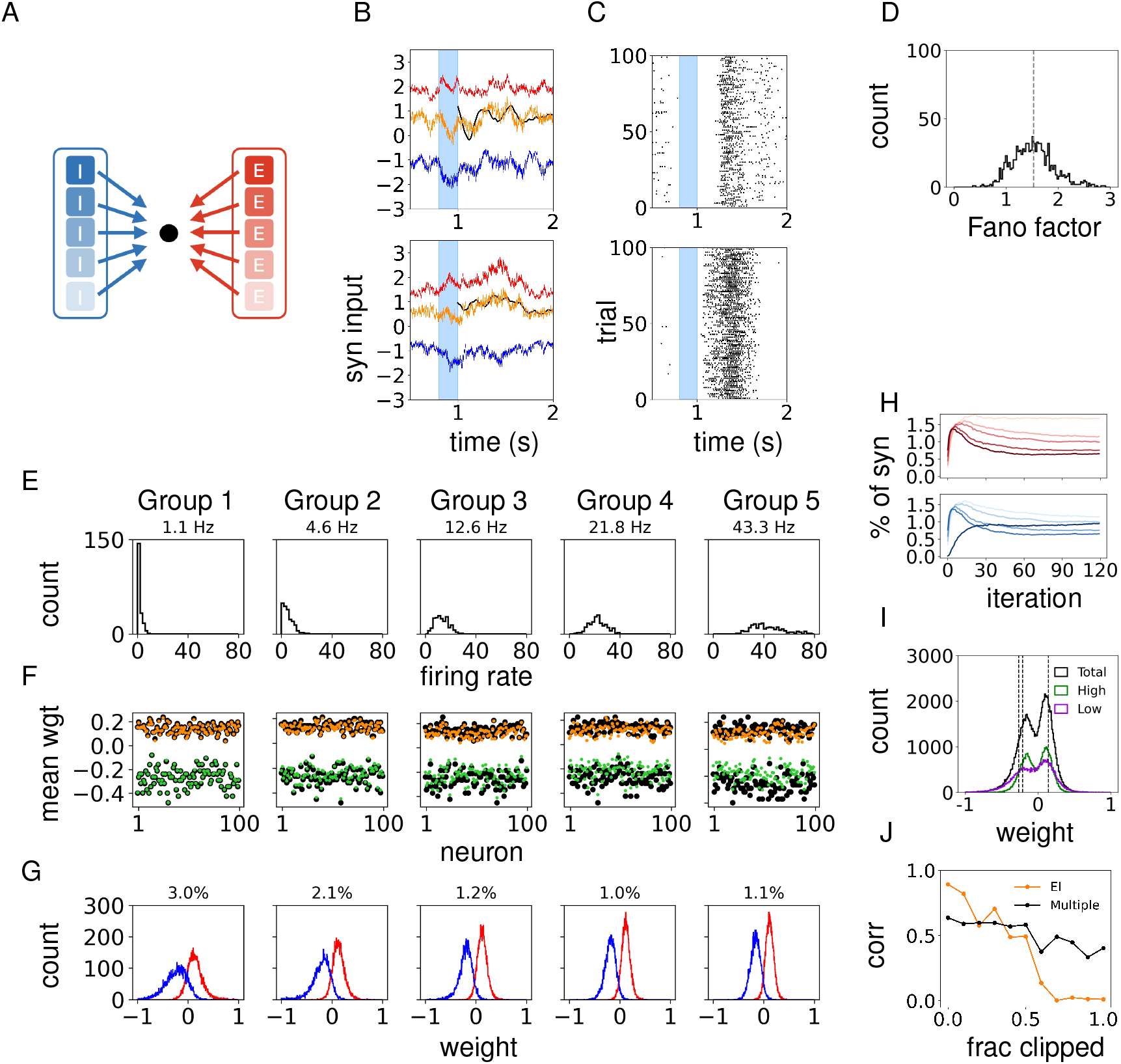
Learning complex activity patterns under the ROWSUM constraint on multiple sub-populations. Target patterns consist of trajectories generated from Ornstein-Ulenbeck process. **(A)** Schematic of constraining multiple subpopulations; black dot indicates one neuron; transparency indicates the within-group firing rate. **(B)** Excitatory (red), inhibitory (blue) and the total (orange) synaptic inputs to trained neurons. **(C)** Spike trains of the same neurons in (a) across multiple trials. **(D)** Fano factor of trained neurons; dashed line indicates the mean. **(E)-(G)** Excitatory and inhibitory neurons are ordered by firing rates and assigned to five subpopulations (i.e., 100 neurons in each group); low to high rate groups are shown from left to right; neurons in the *α^th^* excitatory and inhibitory subpopulations are shown in the same panel, Group *α*. (E) Distribution of firing rates of neurons in Group *α*; the within-group average firing rates shown. (F) Mean strength of synaptic connections that each neuron in the *α^th^* excitatory (orange) and inhibitory (green) subpopulations receive from other neurons in the same subpopulation; black dots show the mean strength of initial connections. (G) Synaptic weight distribution of outgoing connections from neurons in the *α*^*th*^ excitatory (red) and inhibitory (blue) subpopulations. The percentage of synapses (with respect to the total number of synapses in the network) that violates Dale’s law by the neurons in each group are shown. **(H)** Fraction of excitatory (red) and inhibitory (blue) synapses that violates Dale’s law after each training iteration; each line represents outgoing synapses from neurons in the *α^th^* subpopulation; light to dark color shows subpopulation 1 to 5. **(I)** Distribution of trained synaptic weights (black); outgoing synapses from top (green) and bottom (purple) 40% of neurons ordered by firing rates; dashed lines indicate initial synaptic weights. **(J)** Performance of trained networks after removing synapses that violate Dale’s law; constraints on multiple (black) and excitatory-inhibitory (orange) populations.

Overall, we found that the synaptic fluctuations (Fig. 5B) and trial-to-trial spiking variability (Figs. 5C, D; average Fano factor 1.5) increased when multiple sub-populations were constrained, in contrast to the network constrained by the excitatory and inhibitory connections only (Figs. 4B, C).

Next, we combined the *α*^*th*^ excitatory and inhibitory subpopulations into one group and examined their features. The within-group average firing rates increased sequentially in agreement with how neurons were assigned to each group (Fig. 5E; *α*^*th*^ excitatory and inhibitory groups shown in the same panel). The mean strength of synaptic connections from low rate neurons showed little deviations from their initial mean values (Fig. 5F, Group 1), while synaptic connections from high rate neurons (Group 5) showed larger deviations. As will be further analyzed in Results / Section 7, the firing rates of presynaptic neurons and the inverse of the number synapses constrained under the ROWSUM regularization are among the factors that determine how much recursive least squares algorithm can change the mean synaptic weights. In light of this analysis, the high firing rates and the reduced number of neurons in a subpopulation provide an explanation why the mean synaptic weights from neurons in high rate groups deviated from the initial mean weights.

Importantly, the fraction of synapses that violated Dale’s law was reduced (8%; Figs. 5G, H) compared to that found in the network trained under constrained excitatory and inhibitory connections (14%; Fig 4F). Also, the distribution of synaptic weights from the low and high firing rate neurons no longer exhibited drastic differences (Fig. 5I). Interestingly, the fraction of synapses that violated Dale’s law was smaller in groups consisting of high firing rate neurons (see the percentages shown on the panels of Fig. 5G). Tracking the synapses that violated Dale’s law during training revealed that, although such synapses increased rapidly during the early phase of training, they diminished quickly and stabilized to lower values, particularly in the high firing rate groups (Fig. 5H). Moreover, removing the synapses that violated Dale’s law did not significantly affect the performance in networks trained under multiple subpopulation constraints; however, a network trained under the excitatory and inhibitory constraints suffered from reduced performance (Fig. 5J).

### 6 Training initial networks with random connectivity

In this section, we turn to initial networks for which the sums of excitatory and inhibitory synaptic weights to each neuron, respectively, are no longer identical across neurons. We constructed an Erdos-Renyi graph with connection probability *p*, which resulted in a non-constant indegree distribution, and assigned constant weights to each connection type *W*_*αβ*_ for *α, β* ∈ {*E, I*}. To revive the inactive neurons present in such a network, we selected bottom half of the neurons from their mean synaptic weight distribution and added a correction term to the incoming synaptic weights such that the sum matched the expected value *pN*_*β*_*W*_*αβ*_ (See Methods / Section 1 for details). The resulting network exhibited a wide firing rate distribution including many low firing rate neurons (Fig. 6A.i) and large trial-to-trial spiking variability (Figs. 6A.ii-iii).

**Figure 6:**
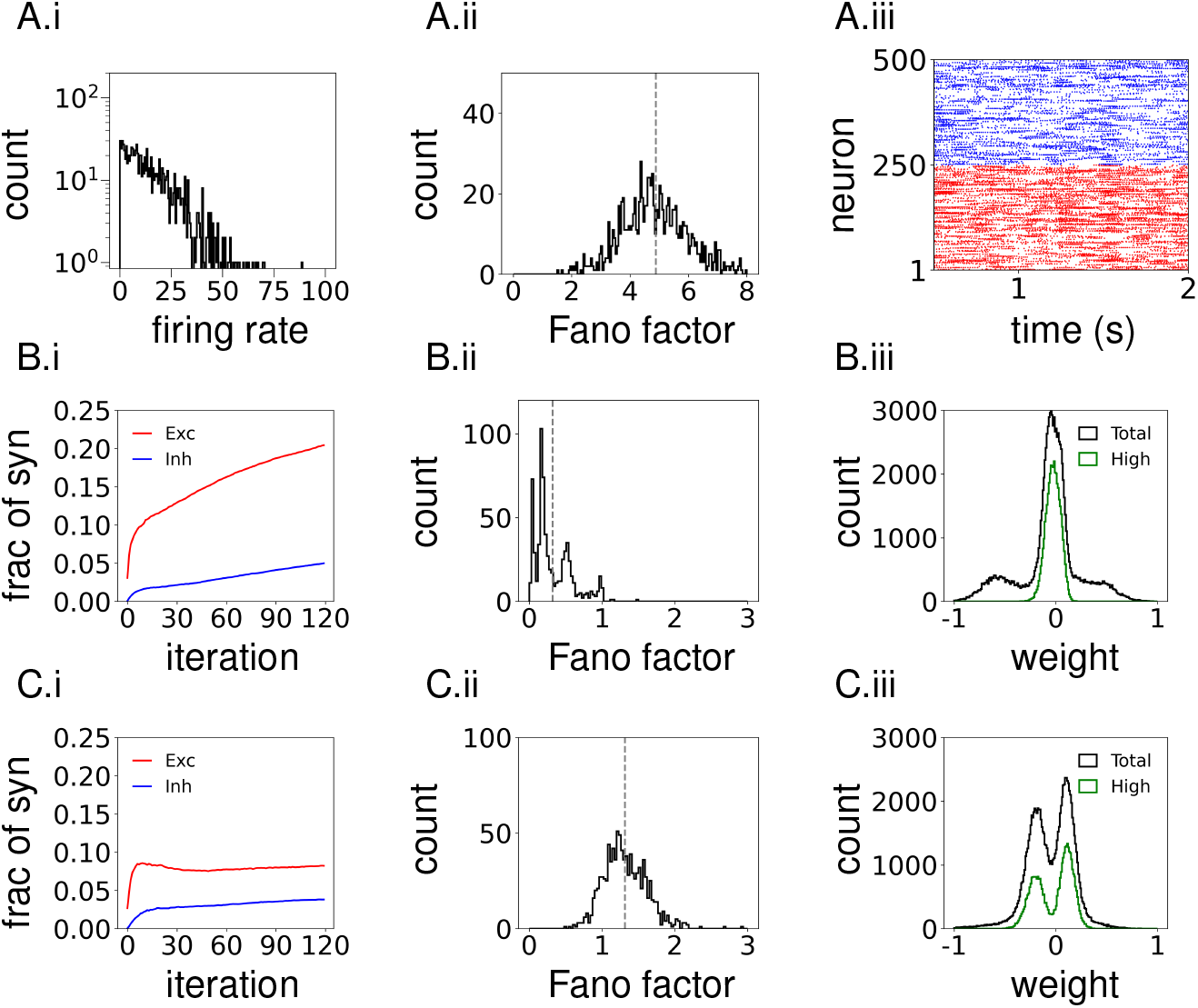
Training initial networks with random connectivity. Synaptic weights to neurons in the bottom half of mean synaptic weight distribution are corrected to their expected values *pN_β_W_αβ_*. Target patterns consist of sinusoidal waves with random phases and fixed amplitude. **(A)** Population activity of an initial network. (A.i) Distribution of neurons’ firing rates. (A.ii) Distribution of neurons’ Fano factor. (A.iii) Spike raster of sample neurons. **(B)** Network trained with the ROWSUM constraint on excitatory and inhibitory synapses. (B.i) Fraction of synapses that violate Dale’s law after each training iteration. (B.ii) Fano factor of trained neurons. (B.iii) Synaptic weight distribution of a trained network (black); Outgoing synapses from top 40% of neurons ordered by firing rates (green). **(C)** Same as in (B), but for a network trained with the ROWSUM constraint on multiple subpopulations.

To avoid compounding the effects of network initialization and target choices, we chose sinusoidal functions as the target patterns since they minimally altered the initial firing rate distribution. Then, we trained the initial network with two different ROWSUM constraints developed in Sections 4 and 5.

First, when only the excitatory and inhibitory synaptic connections to neurons were constrained, the fraction of synapses that violated Dale’s law increased with training iterations (Fig. 6B.i) and the spiking activities of the trained network was highly regular across trials (Fig. 6B.ii). We found that, similarly to networks trained to learn complex target patterns (Fig. 4H), the synaptic connections from high firing rate neurons were degenerating (Fig. 6B.iii).

On the other hand, when the ROWSUM constraint was applied to multiple excitatory and inhibitory subpopulations as discussed in Section. 5 (i.e., five sub-populations for the excitatory and inhibitory neurons, respectively), the fraction of synapses violating Dale’s law remained low (Fig. 6C.i) and the spiking variability was relatively large with the mean Fano factor close to 1.5 (Fig. 6C.ii). Moreover, the synaptic weight distribution, particularly the synaptic connections from high rate neurons, did not converge towards zero (Fig. 6C.iii). These training results demonstrated that applying the ROWSUM constraint on multiple subpopulations can be an effective training scheme even when the the initial network included random connectivity that generated a wide firing rate distribution.

### 7 Analytical upper bound on synaptic weight updates

Since retaining the strong coupling strength of the initial network was critical for generating large synaptic fluctuations and inhibition-dominant dynamics, we further inquired how the ROWSUM-training was able to tightly constrain the mean synaptic weights, such that the mean excitatory and inhibitory synaptic weights to each neuron stayed close to their initial values (Figs. 2B,C, 4E and 5F). In particular, we sought to derive an analytical upper bound on the changes in the sums of excitatory and inhibitory synaptic weights, respectively, in terms of the network and training parameters.

For the analysis, we considered a vector of synaptic connections **w** to an arbitrary neuron *i* in the network whose weights were modified by the RLS algorithm under the ROWSUM-regularization. Elements of **w** are denoted by *w*_*j*_ = *W*_*ij*_ for *j* = 1,…, *K* where *K* is the total number of synapses modified under ROWSUM-regularization. Then, the sums of excitatory and inhibitory synaptic weights are defined, respectively, by

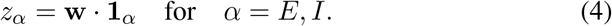

Here, **1**_*E*_ denotes a vector (1,…, 1, 0,…, 0) where the number of 1’s is equal to the number of excitatory synapses *K*_*E*_ modified under ROWSUM-regularization and the rest of *K*_*I*_ = *K* − *K*_*E*_ elements is 0. **1**_*I*_ = (0,…, 0, 1,…, 1) is defined similarly for the inhibitory synapses modified under the ROWSUM-regularization. From the RLS algorithm (Eq. 19 in Methods), the changes in the sums of excitatory and inhibitory synaptic weights at each synaptic update obey

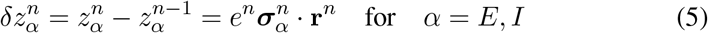

where

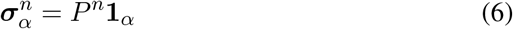

and

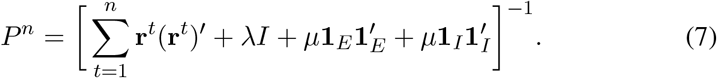

Since 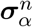 was the only term in the synaptic update rule (Eq. 5) that depended on training and network parameters, it constrained the upper bound on synaptic weight changes. In simulations, the elements of 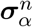 fluctuated in the early phase of training but stabilized as the training iteration *n* increased (Fig. 6A). Based on this observation, we derived an approximation of 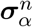 for large *n* and obtained an estimate on its upper bound.

The analysis of 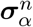 presented below revealed that (1) the initial vector 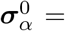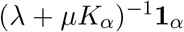 for *α* = *E, I* can be made arbitrarily small if the product *μK*_*α*_ of ROWSUM-penalty *μ* and the number of modified synapses *K*_*α*_ is large (see Methods / Section 5.1), (2) 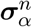 is bounded by 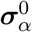 uniformly in *n* ≥ 1 up to a uniform constant (see Methods / Section 5.2), and (3) the update size of mean synaptic weights 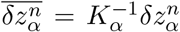 has an upper bound with a scaling factor *O*(1/(*μK*_*α*_)) thanks to (1) and (2) (see Methods / Section 5.3). This analysis result showed that the product of *μ* and *K*_*α*_ tightly constrained the mean synaptic updates because 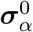 effectively served as a uniform upper bound on 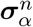 that persisted throughout the training iterations *n*.

An approximation of 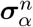 was derived in a few steps. First, the spiking activity **r**^*t*^ during training was approximated by the firing rates 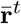 induced by the target patterns and a noise term 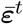 that modeled the spiking fluctuations around the firing rates (see Methods / Section 5.2 for details). Such an approximation was based on the observation that spiking activity stayed close to the target patterns during training due to the fast weight updates (e.g., every 10 ms) by the RLS algorithm.

Next, since the target patterns were learned repeatedly, firing rates 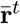 were assumed to have a period equal to the target length (*m*-steps) and repeated *d*-times to cover the entire training duration, i.e., *n* = *d · m*. Assuming such firing rate patterns, the correlation matrix was approximated by 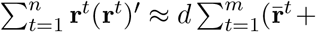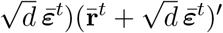 and substituted into the definition of *P* ^*n*^ (Eq. 7) to obtain

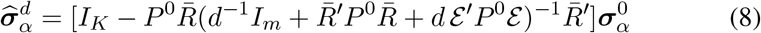

Where 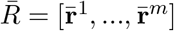 and 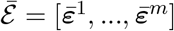. We verified numerically that 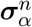 and 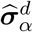 decrease monotonically for large *n* or *d* (FIG.6B), and 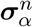 can be estimated by 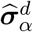 (Fig. 6C). Then, by monotonicity, we can find 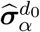 for some constant *d_0_* that serves as a uniform upper bound on 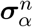:

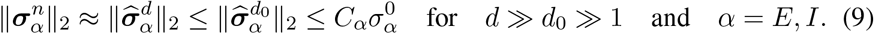

Moreover, the constant *C*_*α*_ is independent of not only *n* but also *μ* and *K*_*α*_ because *P* ^0^ converges elementwise when *μ* or *K*_*α*_ becomes large (see Methods Section 5.3 for details).

Finally, we applied the upper bound on 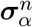 (Eq. 9) to the synaptic learning rule (Eq. 5) to estimate the size of mean synaptic weight updates:

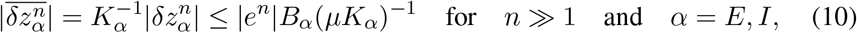

where 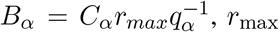 is the maximal firing rate induced by the target patterns, *q*_*α*_ = *K*_*α*_/*K* is the proportion of *α* = *E, I* synapses relative to the total number of synapses *K* modified under ROWSUM-regularization, and *e^n^* is the learning error.

The analytical bound obtained in Eq. 10 suggested two factors that determine the size of synaptic updates. First, the ROWSUM-penalty *μ* constrained the upper bound of 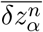 with a scaling factor *μ^−^*^1^. To verify this analytical estimate, we set up an initial network and trained all the synaptic weights with different *μ*-values across training sessions. We evaluated the mean excitatory and inhibitory synaptic weights each neuron received, and then averaged over all neurons in the network. We found that the total changes in the mean excitatory and inhibitory weights after training 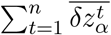 decreased with *μ* consistently to the analytical estimate *O*(*μ^−^*^1^) (Fig. 6D).

Second, the upper bound on the size of mean synaptic weight updates 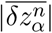 was inversely proportional to *K*_*α*_ if it was sufficiently large and *q*_*α*_ = *K_α_/K* remained constant. In other words, increasing the number of *α* = *E, I* synapses modified under the ROWSUM-regularization effectively strengthened the ROWSUM-penalty. To verify this analytical result, among *pN* synapses each neuron received, a fraction *f* = *K/*(*Np*) of synapses was trained under the ROWSUM-regularization while the rest 1 − *f* of the synapses were trained under the standard *L*_2_-regularization. The fraction *f* of ROWSUM-regularized synapses was increased systematically, while the proportion *q*_*α*_ = *K*_*α*_/*K* of ROWSUM-modified *α* = *E, I* synapses were fixed to 0.5 for both *α* = *E, I*. After training, we evaluated the mean excitatory and inhibitory synaptic weights trained under ROWSUM-regularization and confirmed that the total changes in average synaptic strength decreased consistently with the analytical estimate 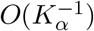 (Fig. 6E).

## Discussion

Recurrent neural networks can learn to perform cognitive tasks (Mante et al., 2013; Sussillo et al., 2015; Chaisangmongkon et al., 2017; Wang et al., 2018) and generate complex activity patterns (Kim and Chow, 2018; Rajan et al., 2016; Laje and Buonomano, 2013). However, due to the large number of connections, the connectivity structure that achieves the learning goal is nonunique (Song et al., 2016; Sussillo et al., 2015). Therefore, it is important to further constrain the recurrent connectivity structure in order to identify solutions relevant to biological neural circuits. In this study, we showed that trained networks can maintain strong excitatory and inhibitory synaptic strength and, consequently, exhibit dynamic features of strongly coupled cortical circuit models, such as stochastic spiking and inhibition-stabilization, if the mean synaptic strength of the strongly coupled initial network is preserved throughout training.

In a related work, Ingrosso and Abbott (2019) developed a training scheme that maintained strong synaptic strength in trained networks by setting up a strongly coupled excitatory-inhibitory network and penalizing the deviation of synaptic weights from their initial values. Specifically, they considered a regularizer 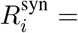 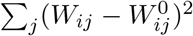 for each neuron *i* that constrained the *L*_2_-distance of individual our work is that the regularizer 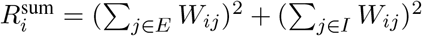 considered in our study constrained the sums of excitatory and inhibitory weights, respectively, but not the individual synaptic weights. In fact, including 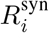 to the cost function is equivalent to adding the standard *L*_2_-regularizer 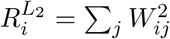 if the synaptic update rule is derived through the RLS algorithm. This is because the effects of subtracting constants 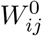 from *W*_*ij*_ are nullified when the gradients at two time points are compared to derive a synaptic update rule (see Methods / Section 3). Bounded coordinate descent algorithm implemented by Ingrosso and Abbott did not suffer from this issue, and they showed that regularizing the weights with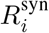, but not 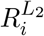, allowed them to preserve the strong synaptic strength of initial networks. For the RLS algorithm, strengthening the *L*_2_-regularizer 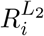 will keep the weights close to their initial values. Our study demonstrated that constraining the mean synaptic strength through 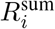 imparts the flexibility for individual synapses to learn the task while maintaining strong coupling of the averaged synaptic weights.

Separation of excitatory and inhibitory neurons is an important feature of biological neural circuits that has implications to the dynamical state of the network. In particular, strong recurrent connections within and between the excitatory and inhibitory neurons give rise to dynamic properties such as asynchronous spiking (van Vreeswijk and Sompolinsky, 1996; Renart et al., 2010; Rosenbaum et al., 2017), balanced amplification (Murphy and Miller, 2009), and inhibition-stabilization (Tsodyks et al., 1997). In recent studies, training schemes were developed to preserve the excitatory-inhibitory structure of recurrent connections. For instance, “clipping” methods skipped synaptic updates that violated the sign constraints (Ingrosso and Abbott, 2019; Kim and Chow, 2018); back-propagation-through-time (BPTT) was adapted to rectify the synaptic weights in order to produce zero gradients in the opposite sign regime (Song et al., 2016); in cases where only the read-out and/or feedback weights were trained, the strong recurrent synapses of the initial network remained unchanged while other weighs were constrained to respect the overall excitatory-inhibitory structure (Nicola and Clopath, 2017). However, these studies focused on respecting the excitatory-inhibitory structure without promoting the strong recurrent connections essential for generating novel network dynamics. Our study suggests that regularizing by imposing additional constraints on the mean synaptic weights can lead to trained networks that inherit dynamic properties of a strongly coupled excitatory-inhibitory network.

In a recent work, Baker et al. (2019) considered network and stimulus configurations that deviated from the standard balanced state in that excessive inhibition present in such a state resulted in threshold-linear stationary rates as a function of external stimulus, as opposed to the linear relationship between positive stationary rates and external inputs in the balanced state. Using the nonlinearity present in threshold-linear activation, they showed that nonlinear computations was possible in the inhibition-excessive excitatory-inhibitory network, pointing out the importance of dynamic regime determined by the synaptic connectivity.

Developing general purpose machine learning algorithms for training spiking neural networks is an active area of research that has made significant progress in the last several years. Recent work has demonstrated that including biological mechanisms, such as spike frequency adaptation, can enhance a spiking neural network’s capability (Bellec et al., 2018), and biologically inspired alternatives to BPTT can be applied to perform supervised and reinforcement learning tasks (Bellec et al., 2019). This progress provides a fertile ground to discover the potential role of connectivity constraints present in biological neural circuits. Spiking neural networks trained with relevant biological features and connectivity constraints may serve as a biologically interpretable alternative to artificial neural networks that can provide further insights into the inner working of biological neural circuits.

## Methods

### 1 Spiking network model

The spiking network model consisted of an equal number of *N*_*E*_ excitatory and *N*_*I*_ inhibitory leaky integrate-and-fire neurons connected randomly with connection probability *p*. The network activity was governed by a set of equations

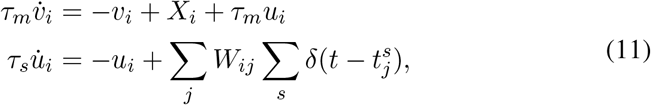

where *v*_*i*_ is membrane potential of neuron *i*, *τ*_*m*_ is membrane time constant, *X_i_* is external input, *u*_*i*_ is synaptic current to neuron *i*, the spike threshold *τ*_*s*_ is synaptic time constant, *W*_*ij*_ is recurrent synaptic weights from neuron *j* to *i* and 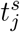 is spike times of neuron *j*. Neurons emitted spikes when the membrane potential reached the spike threshold *v*_*thr*_ = 1, after which the membrane potential was immediately reset to *v*_*reset*_ = 0. Specific parameter values used in simulations are summarized in Table 1.

**Table 1:** Default simulation and training parameters. Any differences from these parameters are mentioned in the main text.

| Neuron parameters |  | Values |
| --- | --- | --- |
| $\tau_m$ | membrane time constant | 10 ms |
| $v_{thr}$ | spike threshold | 1 |
| $v_{reset}$ | voltage reset | 0 |
| Network parameters |  |  |
| $N$ | number of neurons | 1000 |
| $N_E$ | number of excitatory neurons | 500 |
| $N_I$ | number of inhibitory neurons | 500 |
| $p$ | connection probability | 0.1 |
| Synaptic parameters |  |  |
| $\tau_s$ | synaptic time constant | 20 ms |
| $K$ | mean indegree | $pN$ |
| $K_E$ | mean excitatory indegree | $pN_E$ |
| $K_I$ | mean inhibitory indegree | $pN_I$ |
| $W_E$ | excitatory synaptic weight | $1/\sqrt{K_E}$ |
| $W_I$ | inhibitory synaptic weight | $-1.5/\sqrt{K_I}$ |
| $X$ | external input | $0.1\sqrt{K_I}$ |
| $\gamma_E$ | relative strength of recurrent excitation to excitatory population | 1.0 |
| $\gamma_I$ | relative strength of recurrent inhibition to excitatory population | 1.25 |
| $\gamma_X$ | relative strength of external input to excitatory population | 1.5 |
| $W_{EE}$ | $E$ to $E$ synaptic weight | $\gamma_E W_E$ |
| $W_{IE}$ | $E$ to $I$ synaptic weight | $W_E$ |
| $W_{EI}$ | $I$ to $E$ synaptic weight | $\gamma_I W_I$ |
| $W_{II}$ | $I$ to $I$ synaptic weight | $W_I$ |
| $X_E$ | external input to excitatory neurons | $\gamma_X X$ |
| $X_I$ | external input to inhibitory neurons | $X$ |
| Training parameters |  |  |
| $\lambda$ | penalty for $L_2$-regularization | 0.1 |
| $\mu$ | penalty for ROWSUM-regularization | 2.0 |
| $N_{loop}$ | number of training iterations | 120 |
| $T$ | length of target patterns | 1000 ms |

The initial recurrent synaptic weights *W*_*ij*_ and the external inputs *X*_*i*_ were sufficiently strong such that the total excitatory inputs to a neuron, i.e. the sum of excitatory current and external input, exceeded the spike threshold; therefore, feedback inhibition was necessary to stabilize the network activity. Each synaptic weight was determined by its connection type, i.e., *W*_*ij*_ = *W*_*αβ*_ for neuron *j* in population *β* to neuron *i* in population *α*, and neurons in the same population received identical inputs, i.e. *X_i_* = *X*_*α*_ for *i* ∈ *α* where *α, β* ∈ {*E, I*}. The relative strength of recurrent excitation, recurrent inhibition and external input to the excitatory population, i.e., *γ*_*E*_ = *W*_*EE*_/*W*_*IE*_, *γ*_*I*_ = *W*_*EI*_/*W*_*II*_, *γ*_*X*_ = *X*_*E*_/*X*_*I*_, were chosen to be consistent with the balanced condition (van Vreeswijk and Sompolin-sky, 1998; Renart et al., 2010; Rosenbaum and Doiron, 2014):

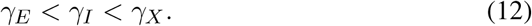

Finally, the recurrent synaptic weights and the external inputs were defined with reference to 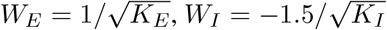 and 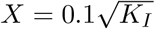:

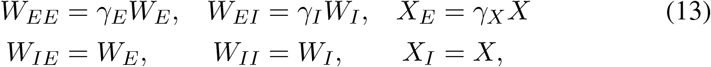

where *K*_*E*_ = *pN*_*E*_ and *K*_*I*_ = *pN*_*I*_ are the mean excitatory and inhibitory indegrees, respectively.

In case of an instantaneous synapse (i.e. *τ*_*s*_ = 0), one spike with weights *W*_*E*_ and *W*_*I*_ would deflect the membrane potential by 0.14 and 0.21, respectively, with the chosen network size *N*_*E*_, *N*_*I*_ = 500 and connection probability *p* = 0.1. Since the spike threshold was *v*_*thr*_ = 1, approximately 8 incoming spikes were enough to trigger a spike. However, due to the synaptic filtering, the actual deflection in membrane potential by a spike was scaled by a factor 1/*τ*_*s*_.

We investigated several types of random connectivity structures for the initial network. First, we considered a network in which the number of synaptic connections to each neuron was fixed to the mean indegree *pN*, but the *pN*_*E*_ excitatory and *pN*_*I*_ inhibitory presynaptic neurons were selected randomly. The synaptic weights in this network structure were set to constants as described in Eq. 13. Since the mean synaptic weights were identical across neurons, we observed homogeneous firing rate distribution in this connectivity type (Fig. 1A.iv). Previous studies considered spiking networks with fixed indegree and constant weights (Brunel, 2000; Ostojic, 2014; Nicola and Clopath, 2017). This network structure was trained on sinusoidal functions in Sections. 1, 2 and 3 and on complex target functions in Sections. 4 and 5.

Second, we constructed an Erdos-Renyi graph where each synaptic connection was generated independently with probability *p*, resulting in a connectivity matrix with a non-constant indegree distribution. The synaptic weights were assigned in two different ways. They were either constants as described in Eq. 13 or were sampled from Gaussian distributions with mean *W*_*αβ*_ and standard deviation *W*_*αβ*_/5 to ensure Dale’s law was not violated. In these networks, the sum of excitatory and inhibitory synaptic weights to each neuron, i.e., Σ_*j* ∈ *E*_ *W*_*ij*_ and Σ_*j*∈*I*_ *W*_*ij*_, respectively, were not identical across neurons, therefore resulted in a wide firing rate distribution including many low firing rate neurons (Fig. 6A.i). To reconcile the differences in synaptic weights across neurons, we added a correction term 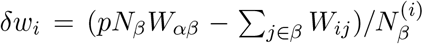 to all nonzero synaptic connections from population *β* to neuron *i* ∈ *α*, where 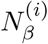 is the number of nonzero connections, such that the total synaptic weight became *pN*_*β*_*W*_*αβ*_. In Section. 2, we considered an Erdos-Renyi connectivity with Gaussian weights in which the synaptic connections to all the neurons were corrected (see Fig. 2C). On the other hand, in Section. 6, we discussed a random connectivity with constant weights in which the synaptic connections were corrected in only a fraction of neurons.

### 2 Training a network of excitatory and inhibitory neurons

Previous studies have shown that the recurrent synaptic connections can be trained using recursive least squares (RLS) to generate target activity patterns (Kim and Chow, 2018; Laje and Buonomano, 2013; DePasquale et al., 2018; Rajan et al., 2016). Specifically, the goal was to train the recurrent connections *W*_*ij*_ such that the synaptic drive *u*_*i*_(*t*) to all neurons *i* = 1,…, *N* in the network learned to follow their own target pattern *f*_*i*_(*t*) defined on time interval [0*, T*] when neurons were stimulated by brief external input. In this study, we included additional penalty terms to the cost function that penalized the sums of excitatory and inhibitory synaptic weights to each neuron, respectively, and then derived the RLS algorithm that included the effects of the additional penalty terms.

The learning rule described below trains the activity of one neuron in the recurrent network. However, the same scheme can be applied to all neurons synchronously (i.e., modify synaptic connections to all neuron at the same time) to update the full connectivity matrix. To simplify notation, we dropped neuron index *i* in the following derivation; **w**= (*W*_*I*1_,…, *W*_*i*K_) denotes a vector of synaptic weights that neuron *i* receives from presynaptic neurons, **r**(*t*) = (*r*_1_(*t*),…, *r*_*K*_(*t*)) denotes the synaptic filtering of spike trains of neurons presynaptic to neuron *i*, and *f* (*t*)*, t* ∈ [0, *T*] is the target pattern of neuron *i*. Neurons were labeled so that the first *K_E_* are excitatory and the remaining *K*_*I*_ = *K* − *K*_*E*_ are inhibitory.

Using linearity in *W*_*ij*_, the equation for synaptic drive *u*_*i*_ was rewritten in the form,

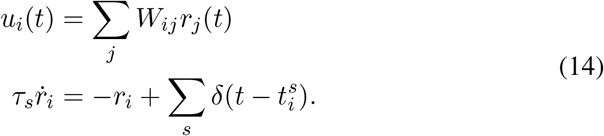

With this formulation, the synaptic drive was expressed as a linear function of connectivity *W*_*ij*_, which allowed us to derive the RLS update rule when the synaptic drive activities were trained.

Here we derive an RLS training algorithm for a network consisting of excitatory and inhibitory neurons applied in Results / Sections 1 to 4. Extension of the ROWSUM constraints to multiple subpopulations are discussed at the end of this section. Since maintaining strong excitatory and inhibitory synaptic strength was critical for generating dynamic features of excitatory-inhibitory network, we constrained the sums of excitatory and inhibitory synaptic weights to each neuron, respectively. The cost function was

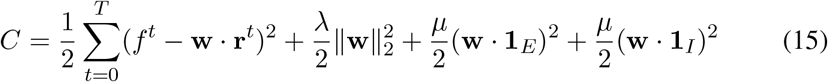

where the first term is the learning error, the second is the *L*_2_-regularization and the remaining two terms comprise the ROWSUM-regularization that penalizes the sums of excitatory and inhibitory synaptic connections to the neuron, respectively. Here 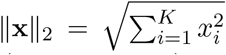 is the standard *L*_2_ norm, **1**_*E*_ = (1,…, 1, 0,…, 0) and **1**_*I*_ = (0,…, 0, 1,…, 1) where the number of 1’s in **1**_*E*_ and **1**_*I*_ is equal to *K*_*E*_ and *K*_*I*_, respectively.

The gradient of the cost function with respect to **w** is

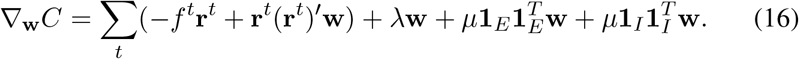

To derive the RLS algorithm, we computed the gradients at two consecutive time points

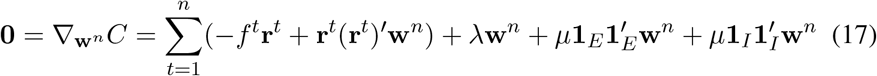

and

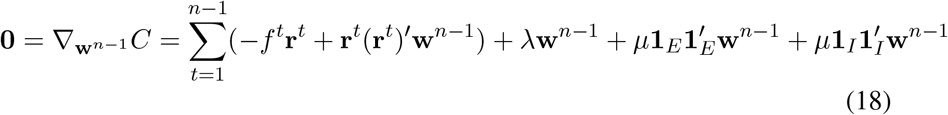

and subtracted Eq. 18 from Eq. 17 to obtain

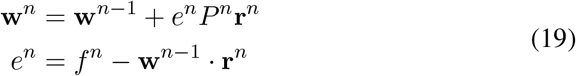

Where

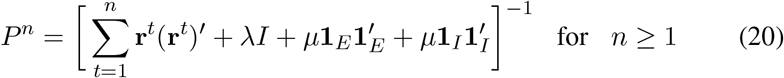

with the initial value

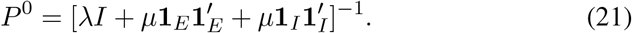

To update *P*^*n*^ iteratively, we used the Woodbury matrix identity

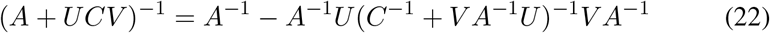

where *A* is invertible and *N*-by-*N*, *U* is *N*-by-*T*, *C* is invertible and *T*-by-*T* and *V* is *T*-by-*N* matrices. Then *P^n^* can be calculated iteratively

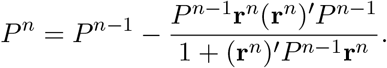

For ROWSUM training on multiple subpopulations, the cost function constrains the sum of synaptic connections from each subpopulation

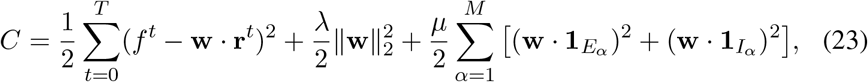

where (**1**_*E*_*α*)_*i*_ = 1 if *i* ∈ *E*_*α*_ and zero otherwise. A sum of multiple block matrices is introduced into the inverse correlation matrix

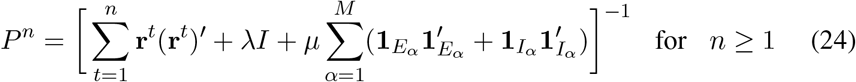

with the initial condition

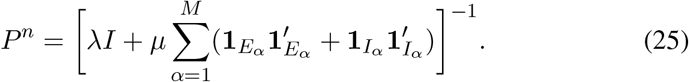

### 3 Translation invariance of RLS algorithm

Here we show that the RLS algorithm is invariant to translation in the sense that adding arbitrary constants to the quadratic penalty terms does not change the synaptic update rule derived from the RLS. Translation invariance arises because of the recursive nature of RLS, i.e., the synaptic update rule uses the difference of the gradient at two time points and thus constants are removed. This justifies the form of the penalty terms in the cost function Eq. 15 since adding a constant to **w** in the regularization terms (e.g., **w** − **m** in the example below) does not affect the synaptic update rule (Eq. 19).

To demonstrate translation invariance, we shifted *w*_*i*_ in the penalty terms by arbitrary number *m*_*i*_ and rewrote the cost function in the following form:

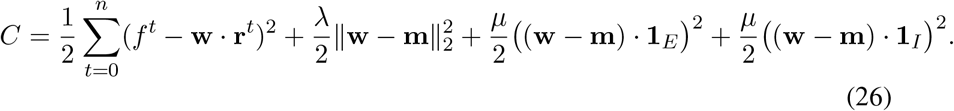

Although this cost function is different from the one considered in the previous section, we now show that the synaptic update rule derived from this alternative form is identical to the previously derived Eq. 19. Differentiating the cost function Eq. 26 with respect to **w**^*n*^ and **w**^*n−*1^ yields

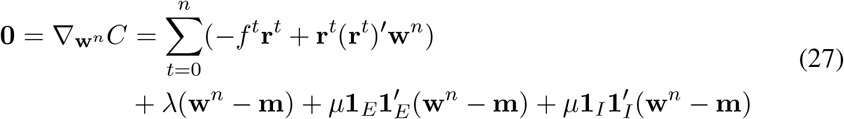

and

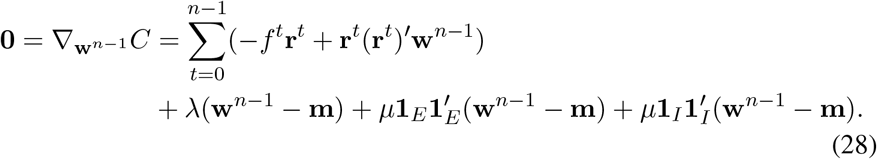

The **m** terms drop out in the final synaptic update rule when we subtract Eq. 28 from Eq. 27. Therefore, the synaptic update rule we obtain from the alternative cost function (Eq. 26) is identical to one (Eq. 19) derived previously from the original cost function (Eq. 15).

### 4 Effects of rate distribution on synaptic weight updates

To gain insights into how the firing rate distribution of presynaptic neurons affects the ROWSUM-training, we derived an analytical expression for the learning rule at the first synaptic update assuming simplified rate distributions. For a homogeneous rate distribution, we considered a rate vector **r**= *r*_0_**1** where all *K* presynaptic neurons have the same rate *r*_0_. For an inhomogeneous rate distribution with high- and low-rate neurons, we considered a rate vector **r**= (*r*_0_,…, *r*_0_*, ϵ,…, ϵ*) where the first *L* neurons fire at rate *r*_0_ and the rest of *K − L* neurons have a small firing rate *ϵ*.

We focused on a ROWSUM constraint applied to connections from all the presynaptic neurons, instead of considering excitatory and inhibitory neurons separately. The first synaptic update can be written as

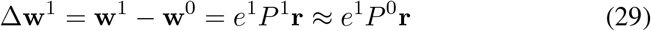

Where

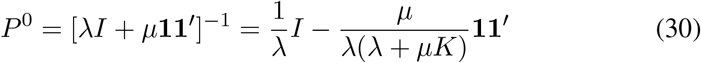

and *P*^1^ = [**r**(**r**)^*I*^ + *λI* + *μ***11**′]^*−*1^ was approximated by *P*^0^ for simplicity.

For the homogeneous rate distribution, the size of synaptic updates is small, i.e. order *O*(1/(*μK*)), for all neurons, thanks to the cancellation of two terms in *P*^0^**1**:

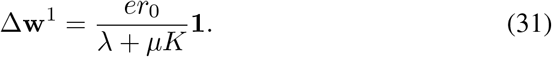

However, for the inhomogeneous rate distribution with high and low rate neurons, the size of the synaptic update can be of order *O*(1) (i.e., note the factors *K* − *L* and *L* in Eq. 32), and the direction of synaptic updates for the high and low rate neurons can be opposite to each other (i.e., note the difference in their signs):

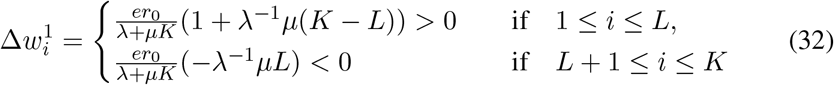

in the limit *ϵ* → 0.

The analysis presented here addresses only the first synaptic update. The full training results can be found in Results / Section 4

### 5 Analytical upper bound on mean synaptic weight updates

In this section, we derived analytical estimates on how the training and network parameters constrained the trained synaptic weights. The sums of excitatory and inhibitory synaptic weights to a neuron were defined, respectively, as

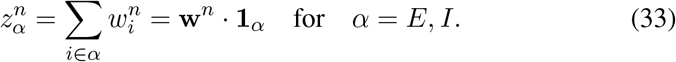

We rearranged the RLS learning rule (Eq. 19) to find an expression for changes in the sums of excitatory and inhibitory synaptic weights:

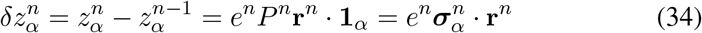

where

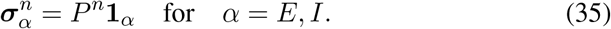

Here we give an overview of the full analysis. We first obtained analytical expressions for the initial value 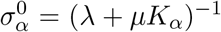 and 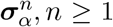 to be used in the subsequent analysis. We found that the initial value 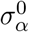 is of order *O*((*μK_α_*)^*−*1^) and used it as the main quantity to bound the rest of 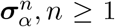 (Section 5.1). To show that 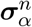 is uniformly bounded in *n*, we decomposed the spiking activity as a sum of spiking rate and noise, and then obtained an approximation 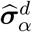 of 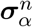 which decreases monotonically and thus has a uniform upper bound 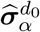 for some *d* ≫ *d*_0_ (Section 5.2). We numerically verified that 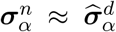, and they both decrease monotonically which led to the uniform upper bound on 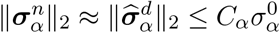 and mean synaptic weights 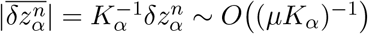 (section 5.3).

The main result deduced from the analysis is that the changes in mean synaptic Weights 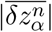 is constrained by the product *μK*_*α*_. By either increasing the *μ*-penalty or the number of synapses *K*_*α*_ modified by the ROWSUM-regularization, the changes in mean synaptic weights can be constrained tightly.

#### 5.1 Analytical expressions for 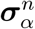

To find the initial 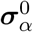 for *α* = *E, I*, we substituted *A* = *λI*_*K*_, *U* = [**1_E_**, **1_I_**], *V* = *U*′ and *C* = *I*_2_ to the Woodbury matrix identity (Eq. 22) to obtain

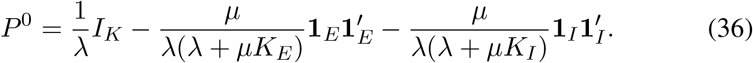

Here the subscript in *I*_*K*_ denotes the dimensionality of the *K* × *K* identity matrix. Then,

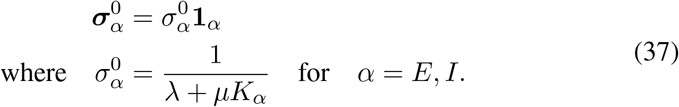

We can see immediately that the initial value 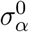 is of order *O*(1/(*μK*_*α*_)). To obtain estimates for 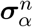 with *n ≥* 1, we used the Woodbury matrix identity (Eq. 22) to express *P ^n^* in the form

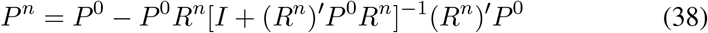

where *R*^*n*^ = [**r**^1^,…, **r**^*n*^] and obtained

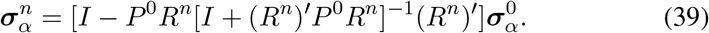

#### 5.2 Asymptotic approximation of 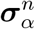

To show that 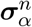 can be bounded uniformly in *n*, it suffices to show that the right hand side of Eq. 39 does not depend on *n*. To this end, we analyzed its asymptotic behavior as the training iteration *n* grew arbitrarily large. We used the fact that (1) the same target pattern was learned repeatedly during training and (2) fast synaptic weight updates (e.g., every 10 ms) by the RLS algorithm kept the spiking activity close to the target pattern. Then, the actual spiking activity **r**^*t*^ was approximated with a firing rate function 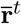 induced by the target patterns where 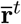 is a periodic function with a period equal to the target length (*m*-steps) and repeated *d*-times to cover the entire training duration, i.e., *n* = *d · m*.

To make it clear that 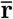 is periodic in time, we used a time index 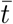 that denotes the time point within the target pattern that have period *m*. Specifically, the spiking activity is expressed as

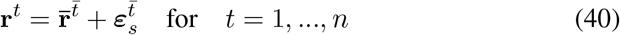

where 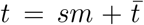 and *s* = 1,…, *d* indicate the training iteration number. We introduced a noise term 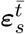 whose *i*^th^ component is the fluctuation of neuron *i*’s spiking activity around the firing rate 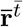 at the *s*^th^ training session. For the following analysis, we assumed a simple noise model that 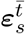 is a multivariate random variable at each 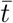 with no correlations between neurons within and across training sessions

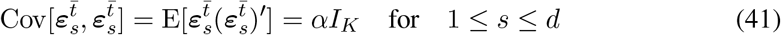

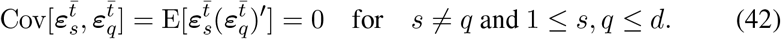

To derive an alternative expression for 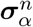 amenable to asymptotic analysis, we expressed the correlation of spiking activities in terms of the firing rates and fluctuations around them:

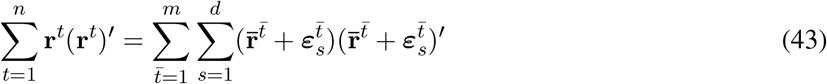

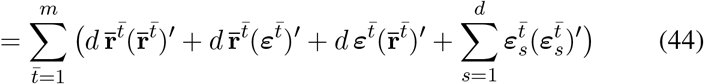

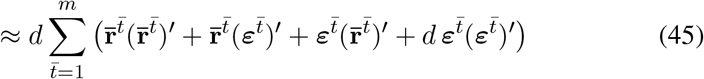

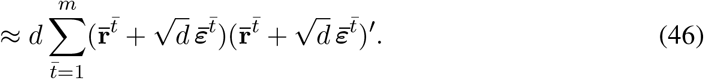

The noise approximation in Eq. 45 is equivalent to following relationship

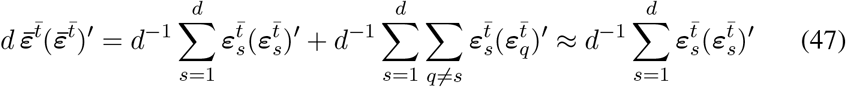

Where

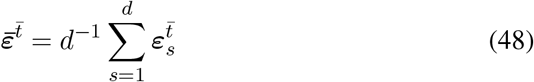

and the cross-training correlations in Eq. 47 was ignored by the noise assumption given in Eq. 42. In Eqs. 45 and 46, the dominant term is 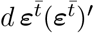, so 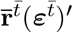 and 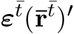 in Eq. 45 were replaced by 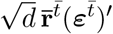 and 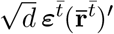 in Eq. 46.

Next, we substituted the approximated Σ*t* **r**^*t*^(**r**)′ (Eq. 46) into the definition of *P*^*n*^ (Eq. 38) and suppressed *O*(*d*)-terms to obtain

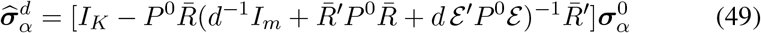

where

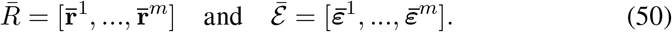

Note that for uncorrelated noise, the term ɛ′*P* ^0^ɛ yields an identity matrix which effectively serves as a regularizer that makes it possible to invert 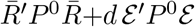.

We verified numerically that the actual 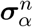 can be well approximated by 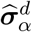 (Fig. 7B). Moreover, since 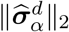 decreases monotonically with *d*, there exists a constant *d*_0_ such that

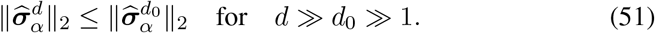

**Figure 7:**
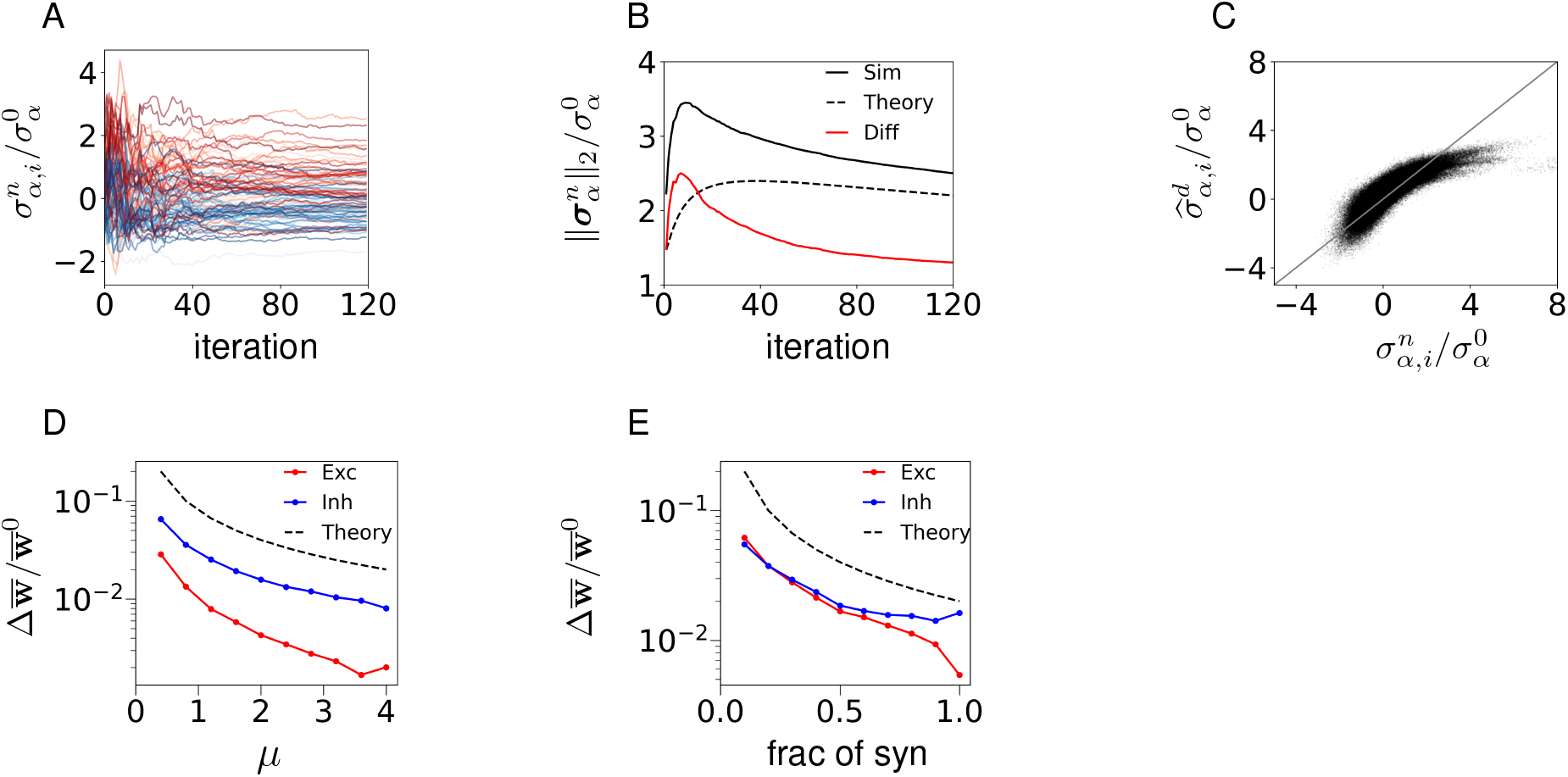
Analytical upper bound on synaptic updates. **(A)** Elements in 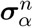 stabilizes if the training iteration *d* becomes large. 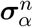 of sample neurons are shown; excitatory (red) and inhibitory (blue). 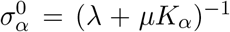 denotes the initial value. **(B)** Comparison of the actual 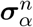 and the theoretical estimate 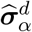 as a function of training iteration *d*; actual (solid), theory (dotted), difference of actual and theoretical (red). **(C)** Comparison of the actual values 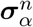 and the theo-retical estimates 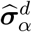 as training iteraton *d* = 120. Scatter plot shows the vector elements of all neurons; identity line (gray). **(D)** Total changes in the mean synaptic weights 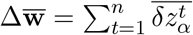 due to training, normalized by the mean initial weight 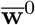, are shown as a function of *μ*; theoretical estimate of the scaling factor *O*(*μ^−^*^1^) (black dotted line); mean excitatory (red) and inhibitory (blue) weights. **(E)** Same as in (D) but varying the number of synapses constrained by ROWSUM regularization; fraction of synapses equals 1 if all synapses are regularized by the ROWSUM constraint. Color code same as (D); theoretical estimate of scaling factor is 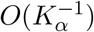.

#### 5.3 Upper bound on the size of synaptic updates

By its definition in Eq. 49, 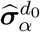 can be expressed in terms of the initial value 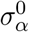

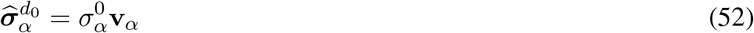

where

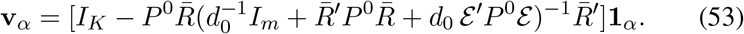

Next we made two observations regarding the upper bound on **v**_*α*_. First, it can be bounded by a constant *C*_*α*_ that does not depend on *d* or *n* since *P*^0^ = *P*^0^(*μ, λ, K*) is a function of the penalty terms *μ, λ* and the number of synapses *K*, and 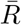 and 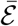 are functions of spiking rate 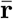 and noise 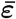, respectively, thus

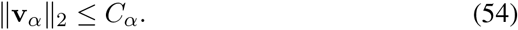

Second, the constant *C*_*α*_ does not depend on *μ* and *K*_*α*_ if they become sufficiently large. This can be seen from the fact that *P*^0^ converged elementwise as *μ* or *K*_*α*_ become arbitrarily large. Specifically,

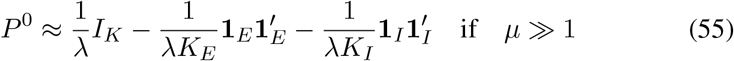

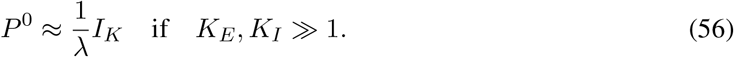

These observations led to the uniform upper bound

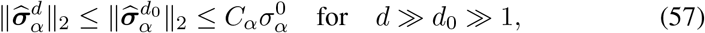

where *C*_*α*_ does not depend on *d, μ* and *K*_*α*_.

Applying this upper bound to the synaptic learning rule (Eq. 34), we obtained that the update size of the sums of excitatory and inhibitory synaptic weights, respectively, at each update are constrained by

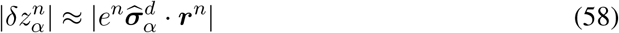

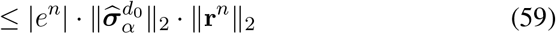

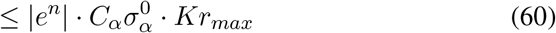

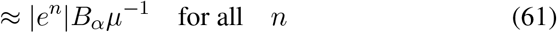

where *B*_*α*_ = *C*_*α*_*r*_*max*_*K/K*_*α*_. We used 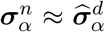 to approximate 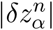 in the first line, the uniform upper bound on 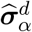 (Eq. 57) in the second and third lines, and 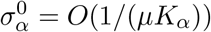 (Eq. 37) in the last line.

This estimate showed that the upper bound on the sum of synaptic weights (1) is inversely proportional to *μ* and (2) does not depend on the number of synaptic connections *K*_*α*_ modified under the ROWSUM-regularization as long as the proportion *q*_*α*_ = *K*_*α*_/*K* remains constant. Finally, we concluded that the upper bound on the average synaptic weights was inversely proportional to the product *μK*_*α*_:

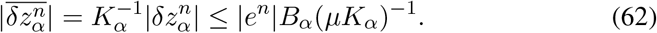

## Acknowledgments

This work was supported by the Intramural Program of the NIH, NIDDK.

